# The SARS-CoV-2 receptor-binding domain facilitates neutrophil transepithelial migration and nanoparticle uptake in the mice airways

**DOI:** 10.1101/2022.04.12.488042

**Authors:** Elena L. Bolkhovitina, Julia D. Vavilova, Andrey O. Bogorodskiy, Yuliya A. Zagryadskaya, Ivan S. Okhrimenko, Alexander M. Sapozhnikov, Valentin I. Borshchevskiy, Marina A. Shevchenko

## Abstract

SARS-CoV-2-induced infection is still dangerous. Mouse models are convenient to the investigation of virus-activated immune response mechanisms. However, mice are not proper model organisms to study COVID-19 due to decreased interaction affinity between the SARS-CoV-2 receptor-binding domain (RBD) and mouse angiotensin-converting enzyme 2 (ACE2) compared with human ACE2. In the present study, we propose a mouse model that allows estimating the influence of SARS-CoV-2 on the immune system. To mimic the effects of RBD– ACE2 high-affinity interaction, mice received the ACE2 inhibitor MLN-4760. To simulate virus loading, we applied 100 nm particles suspended in the solution of RBD via the oropharyngeal route to mice. In this model, MLN-4760 application enhanced neutrophil egress from the bone marrow to the bloodstream and RBD attracted neutrophils to the luminal side of the conducting airway epithelium. By contrast, inert 100 nm particles were not potent to stimulate neutrophil recruitment to the conducting airway mucosa. Using this model, and by altering the dosage of the ACE2 inhibitor, nanoparticles, and RBD, one can adapt it to investigate different COVID-19 states characterized with mild or severe airway inflammation.

**Statement:** This study presents a mouse model that allows estimating the influence of SARS-CoV-2 on the immune system and investigates immune cell-model virus particle interactions in the conducting airway mucosa.

## 3.2.4. Introduction

The interaction of angiotensin-converting enzyme 2 (ACE2) and receptor-binding domain (RBD) of spike protein of SARS-CoV-2 plays an important role in the virus entry into the human organism (Wang et al., 2020). Since SARS-CoV2 is not able to use mouse ACE2 as an entry receptor (Zhou et al., 2020), transgenic mice, which express human ACE2, were developed and used to investigate infection mechanisms (Bao et al., 2020; Winkler et al., 2022). Another approach for mouse model generation uses virus adaptation through several passages of a clinical isolate of SARS-CoV-2 in the respiratory tract of old mice (Gu et al., 2020). Such an approach resulted in a mutation in the RBD domain of the virus, confirming the importance of RBD–ACE2 high-affinity interaction for the infection.

In addition to its role in the virus entry, ACE2 has other physiological functions that can also be implicated in SARS-CoV-2-induced respiratory inflammation (Rossi et al., 2020). The studies of immune and renin-angiotensin systems interplay demonstrate that ACE2 inhibition and subsequent elevation of extracellular Angiotensin II (AngII) level activate neutrophil-mediated inflammatory response (Bernstein et al., 2018; Zhao et al., 2019). The neutrophil-mediated immune response is intrinsic in acute lung injury and observed during severe COVID19 (Cicko et al., 2018; Gusarova et al., 2021; Yang et al., 2021). Neutrophils are known to play a pathogenic role in SARS-CoV-2 induced inflammation, in particular, neutrophil extracellular traps (NETs) were identified in the lungs of patients with severe inflammation (Barnes et al., 2020; Middleton et al., 2020; Veras et al., 2020; Zuo et al., 2020). The formation of NETs is a result of an uncontrolled neutrophil-mediated immune response and contacts between neutrophils and SARS-CoV-2 virus particles (Mutua and Gershwin, 2021; Veras et al., 2020). There have been several hypotheses of the uncontrolled neutrophil-mediated immune response induction by SARS-CoV-2 (Hazeldine et al., 2021). Yet, the mechanisms are still not completely understood.

In this study, using immunohistochemistry and three-dimensional imaging of the whole-mount conducting airways, we investigated the ability of 100 nm fluorescent particles to attract neutrophils after oropharyngeal application to mice. We checked the effects of RBD and the ACE2 inhibition on the neutrophil recruitment and nanoparticle internalization. This study presents a model that mimics some aspects of COVID-19 and affords to investigate the effects of SARS-CoV-2 on the immune system without using humanized mice and virulent pathogens.

## 3.2.5. Results

### RBD induces neutrophil migration through the conducting airway epithelial barrier in response to inhaled nanoparticles

To investigate the conducting airway immune response to SARS-CoV2, mice received an oropharyngeal application of 100 nm particles – FluoSpheres, comparable in size with SARS-CoV2 virus particles (Bar-On et al., 2020). Before application to mice, FluoSpheres were resuspended either in PBS or RBD solution. In both cases, 24 h after the application, FluoSpheres were detected on the luminal side of the conducting airway epithelium (Fig. 1A, B). In both cases, FluoSpheres were mostly internalized by CD169^+^ actin-rich cells; however, free FluoSpheres were also detected. The application of FluoSpheres suspended in PBS did not induce measurable neutrophil migration to the conducting airway mucosa. Both anti-Ly6G and anti-CD11b antibodies revealed that Ly6G^+^ and CD11b^+^ cells were mostly located beneath the smooth muscle layer in the submucosal compartment (Fig. S1A–C). Thus, 24 h after FluoSpheres application, only a few CD11b^+^ were observed in the conducting airway wall, between the epithelium and the smooth muscle layer (Fig. 1A). No CD11b^+^ cells were detected in the luminal side of the epithelium close to FluoSpheres (Fig. 1A). Application of FluoSpheres suspended in RBD solution induced CD11b^+^ cells migration to the conducting airway mucosa and specifically to the luminal side of the epithelium (Fig. 1B). Moreover, migrating through the epithelial barrier CD11b^+^ cells were detected in close proximity to FluoSpheres (Fig. 1B,C, arrows). The precise analysis showed that CD11b^+^ cells together with CD169^+^ cells participated in the internalization of FluoSpheres in the luminal side of the epithelium (Fig. 1D). When FluoSpheres were applied to mice in BSA solution, they did not induce CD11b^+^ cell infiltration to the conducting airway mucosa (Fig. S2B). The RBD solution alone (without FluoSpheres) did not induce neutrophil recruitment to the conducting airway mucosa (Fig. S2C).

**Figure 1.**
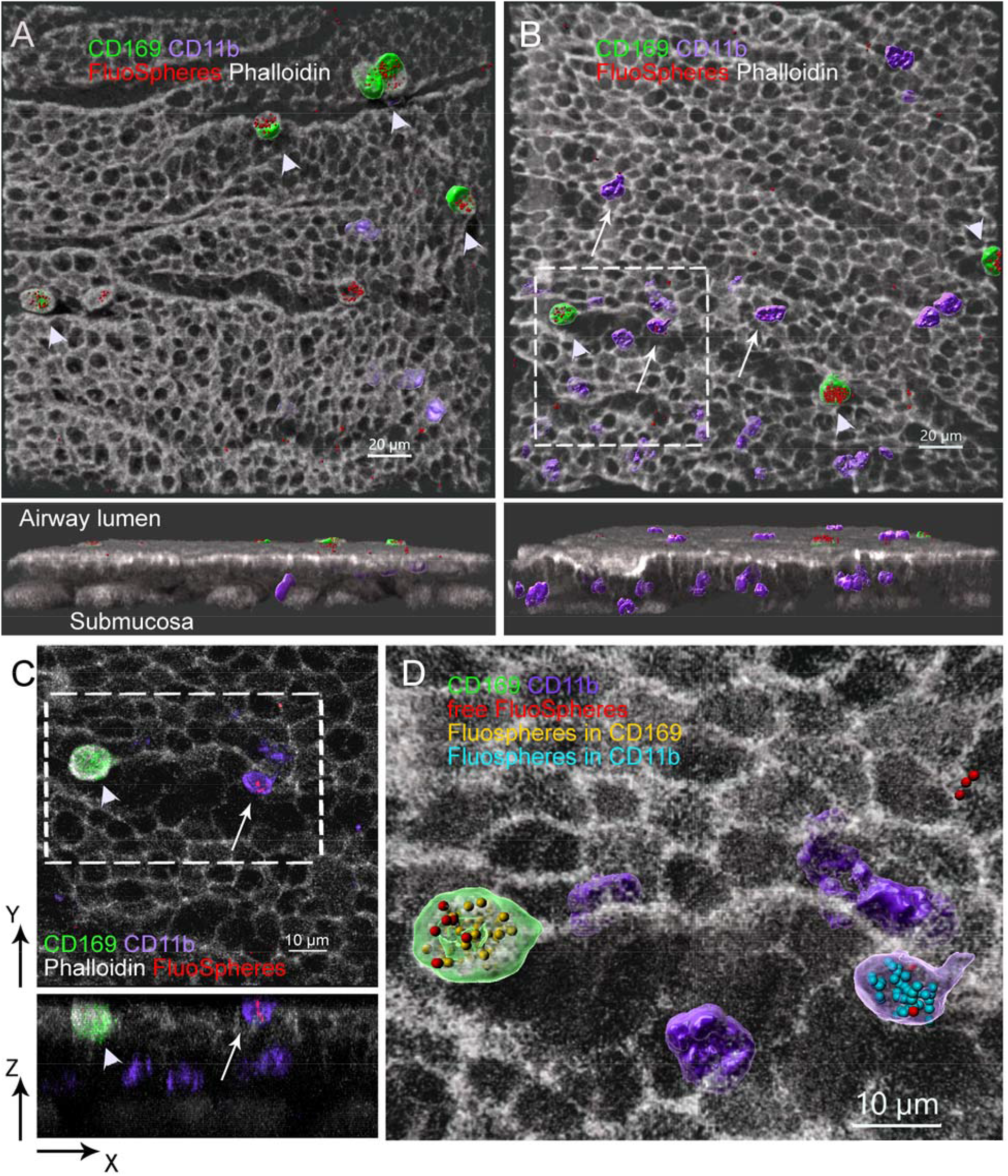
Neutrophils migrate to the conducting airway mucosa of mice in response to fluorescent 100 nm particles suspended in RBD solution. A, B. Representative images of the regions of the conducting airway mucosa of mice at 24 h after the oropharyngeal application of FluoSpheres suspended in PBS (A) or RBD solution (B). Frontal views (upper images) and side views (lower images). FluoSpheres (red), CD169^+^ cells (green), CD11b^+^ cells (violet) are represented via surface rendering, epithelium, and smooth muscles (grayscale) are shown as shadow projection. CD169^+^ cells interacting with FluoSpheres are indicated with arrowheads, CD11b^+^ cells interacting with FluoSpheres are indicated with arrows. Scale bar 20 µm. C. XY (upper image) and XZ (lower image) projections of the region approximately indicated in (B) demonstrate CD169^+^ cell (green), CD11b^+^ cells (violet), FluoSpheres (red), and actin filaments of epithelium and smooth muscles (grayscale) via volume rendering. Scale bar 10 µm. D. Enlarged image of the region boxed in C showing the internalization of FluoSpheres by CD169^+^ cell and CD11b^+^ cell. CD169^+^ cell (green), CD11b^+^ cell (violet), free FluoSpheres (red), FluoSpheres inside or in contact with CD169^+^ cell (yellow) and FluoSpheres inside or in contact with CD11b^+^ cell (cyan) are represented via surface rendering. The FluoSphere surface was enlarged to make it visible. The surfaces of cells that internalized FluoSpheres are transparent to make FluoSpheres inside the cells visible. Scale bar 10 µm.

Previously, we demonstrated that upon inflammation, in the conducting airway mucosa, CD11b^+^ cells were mostly represented by the Ly6G^+^ neutrophils (Bogorodskiy et al., 2020). Thus, our present data support the evidence that FluoSpheres suspended in RBD solution (rather than a phosphate buffer) can attract neutrophils to the conducting airway mucosa and facilitate the ingestion activity of these neutrophils.

### Systemic blocking of ACE2 is necessary for neutrophil recruitment to the bloodstream

Although we demonstrated the RBD contribution to the neutrophil attraction to the conducting airway mucosa, the effect was not caused by the RBD–ACE2 interaction due to low affinity of mouse ACE2 to SARS-CoV-2 RBD. To investigate the influence of systemic ACE2 inhibition on the neutrophil-mediated response, we injected mice with selective ACE2 inhibitor MLN-4760, which was shown to affect mouse ACE2 (Ye et al., 2012). Using the strategy suggested by Liu et al. (Liu et al., 2020) with slight modifications, we estimated the myeloid cell and neutrophil proportions in the periphery blood of mice that received MLN-4760 and control mice (Fig. 2A,B). We showed that a single MLN-4760 intraperitoneal (i.p.) injection significantly increased the blood myeloid cell and neutrophil proportions 2 h after the application (Fig. 2C,D). As the injection itself (as a trauma) can alter the blood neutrophil number, the control group of mice received a PBS i.p. injection. Two hours after, the blood neutrophil proportion in mice that received the PBS injection was significantly higher than that of the intact mice but significantly lower than that of mice injected with MLN-4760 (Fig. 2C,D).

**Figure 2.**
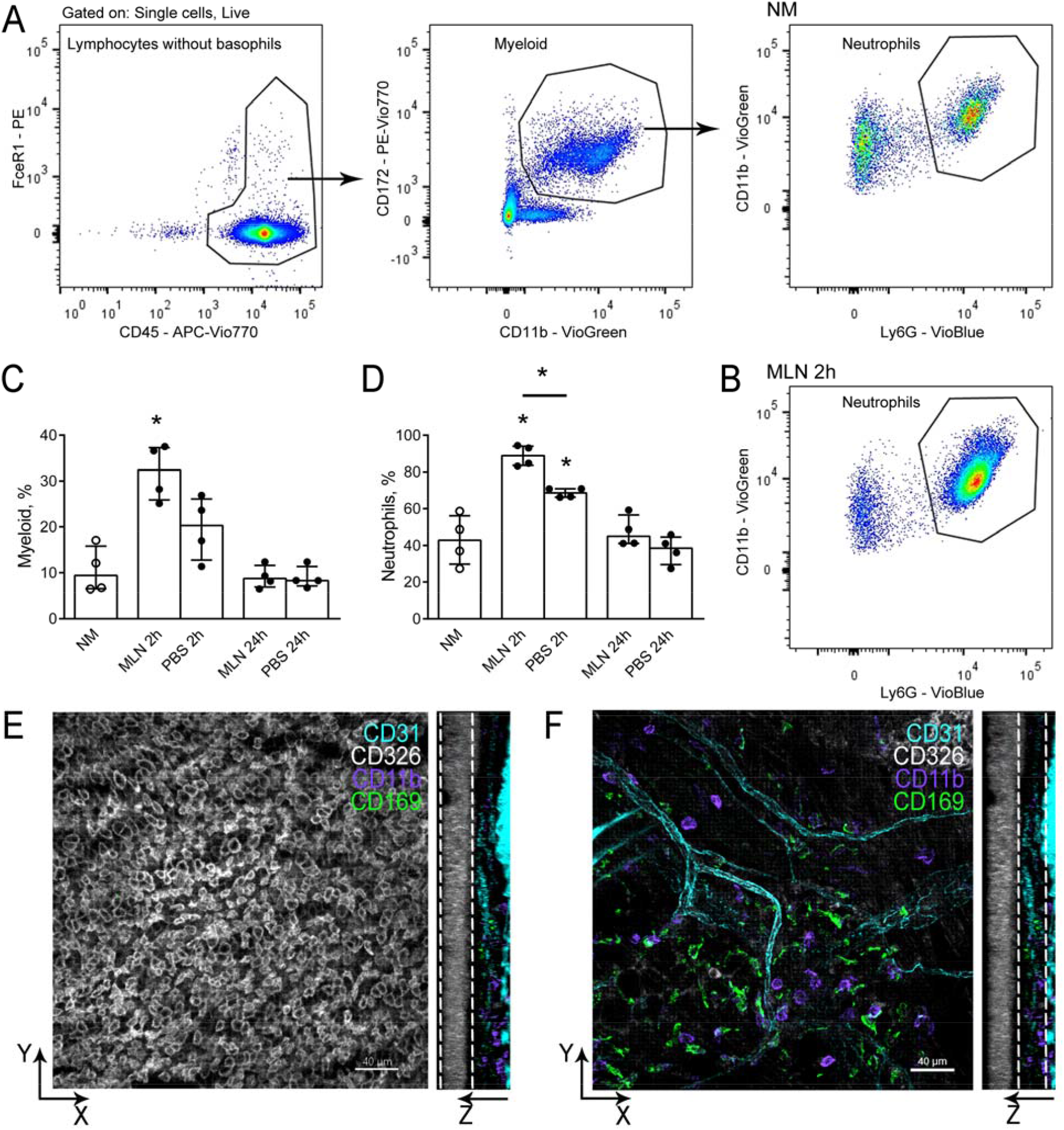
Systemic blocking of ACE2 induces fast neutrophil recruitment to the bloodstream. A, B. Representative flow cytometry plots showing strategy for the neutrophil population identification in the periphery blood of the intact mouse (A) or the MLN-4760-injected mouse (B). C, D. Percentage of the myeloid cells (C) and the neutrophils (D) in the periphery blood of the intact mice (NM), 2 h after the MLN-4760 injection (MLN 2h), 24 h after the MLN-4760 injection (MLN 24h), 2 and 24 h after the PBS injection (PBS 2h and PBS 24h, consequently). The data are pooled from two independent experiments, the total number of mice n=4, and shown as median and i.q.r. Significant difference between the indicated group and the intact mice or between the indicated groups was detected using Mann–Whitney test *: p ≤ 0.05. E, F. Representative images of the regions of the conducting airway mucosa and submucosal compartment of mice 24 h after the i.p. MLN-4760 application. The epithelium (grayscale), blood vessels (cyan), neutrophils (violet), CD169^+^ cells (green) are represented as frontal (X-Y) and lateral (Y-Z) projections. Each frontal projection contains an extended view comprising several Z-sections. The dotted lines approximately indicate the conducting airway mucosa Z-sections that are represented on the frontal projection (E) or the submucosal compartment (F). Scale bar 40 µm.

As expected, the i.p. injection of MLN-4760 did not influence the CD11b^+^ cell migration to the conducting airway mucosa (Fig. 2E) but it affected the neutrophil migration to the lungs. Thus, 24 h after the MLN-4760 i.p. injection, CD11b^+^ cells were detected in the submucosal compartment of the conducting airways of mice (Fig. 2F, Fig. S3A). However, a few CD11b^+^ cells were also identified in the lung tissues of the intact mice (Fig. S3B). Due to the absence of the physical border between the submucosa and the alveolar compartment, quantitative analysis of submucosal CD11b^+^ cells is complicated.

Thus, the i.p. injection of MLN-4760 induces neutrophil recruitment to the blood shortly after the application and does not induce neutrophil accumulation in the conducting airway mucosa without any additional stimuli.

### Blocking of ACE2 in the airways induces the neutrophils migration to the conducting airway mucosa

To estimate the effect of the local airway blocking of ACE2, we applied MLN-4760 to mice via the oropharyngeal route. Similarly to the i.p. MLN-4760 injection (Fig. 2 C, D), we observed a significant elevation of the blood myeloid cell and neutrophil proportions 2 h after the o.ph. application of MLN-4760 (Fig. 3A,B). The o.ph. application of PBS did not alter the myeloid cell and the neutrophil proportions significantly (Fig. 3A,B). Twenty-four hours after both MLN-4760 and PBS application, there were no significant alterations in the myeloid cell and the neutrophil proportions in the blood (Fig. 3B). However, at this time point, in mice that received o.ph. MLN-4760 we detected CD11b^+^ cells in the conducting airway mucosa, particularly, in the conducting airway wall – between the smooth muscle and epithelial layers (Fig. 3C). In mice that received o.ph. PBS, we detected CD11b^+^ cells only beneath the smooth muscles (in the submucosal compartment) (Fig. 3D).

**Figure 3.**
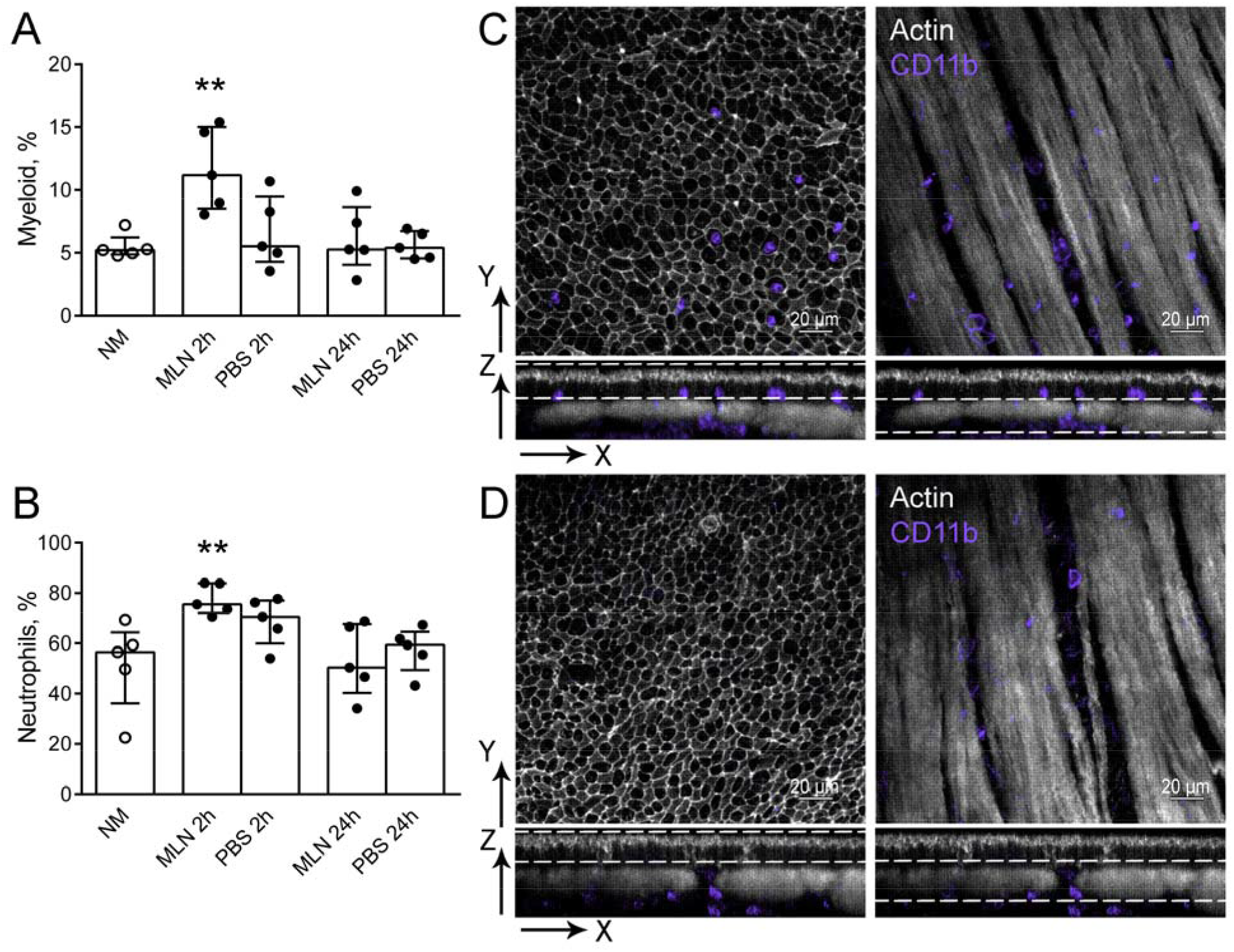
Effects of the oropharyngeal MLN-4760 application to mice on neutrophil recruitment to the bloodstream and the airways. A, B. Percentage of myeloid cells (A) and neutrophils (B) in peripheral blood of intact mice (NM), 2 h after the MLN-4760 o.ph. application (MLN 2h), 24 h after the MLN-4760 o.ph. application (MLN 24h), two and 24 hours after the PBS o.ph. application (PBS 2h and PBS 24h, respectively). The data are pooled from two independent experiments, the total number of mice n=5, and shown as median and i.q.r. Significant difference between the indicated group and the intact mice was detected using Mann–Whitney test **: p ≤ 0.01. C, D. Representative images of the regions of mouse conducting airway mucosa: the epithelium (left images) and the smooth muscle layers (right images), 24 h after the o.ph. application of MLN-4760 (C) or PBS (D). The epithelium (grayscale), neutrophils (violet) are represented as frontal (X-Y, upper images) and lateral (Y-Z, lower images) projections. Each frontal projection contains an extended view comprising several Z-sections. The dotted lines approximately indicate the conducting airway mucosa Z-sections that are represented on the frontal projections. Scale bar 20 µm.

Thus, the local airway ACE2 inhibition elevates the myeloid cell and the neutrophil proportions in the blood in 2 h and attracts CD11b^+^ cells to the conducting airway mucosa in 24 h. The airway ACE2 blocking is not sufficient to attract CD11b^+^ cells to the luminal side of the epithelium.

### Synergetic effect of ACE2 blocking and FluoSpheres application induces neutrophil migration from the blood and retention in the airways

As the MLN-4760 o.ph. application alone stimulated the CD11b^+^ cell migration to the conducting airway mucosa, we checked whether these CD11b^+^ cells can sense FluoSpheres. Firstly, we estimated the synergetic effect of MLN-4760 and FluoSpheres on the systemic neutrophil-mediated response. Similar to the o.ph. application of MLN-4760 alone, the o.ph. application of MLN-4760 together with FluoSpheres significantly elevated the blood myeloid cell and the neutrophil proportions 2 h after the application compared to the intact mice (Fig. 4A,C,D). Interestingly, 24 h after the application, the neutrophil percentage in mice that received the o.ph. application of MLN-4760 and FluoSpheres was significantly lower than that of the intact mice (Fig. 4B-D). At the same time, a measurable amount of CD11b^+^ cells was detected in the lung tissues of mice that received MLN-4760 and FluoSpheres (Fig. 4E). To compare the number of CD11b^+^ cells in the submucosa of mice that received MLN-4760 together with FluoSpheres and MLN-4760 alone, we estimated the number of CD11b^+^ cells that were in contact or in close proximity to the smooth muscles (Fig. 4E, right image, Fig. 4E). Such an approach permitted us to avoid incorrect quantification of the cells from the alveolar space located close to the submucosal cells. The quantitative analysis demonstrated the elevated number of the submucosal CD11b^+^ cells in the conducting airways of mice treated with both MLN-4760 and FluoSpheres compared to the MLN-4760 treatment alone (Fig. 4G).

**Figure 4.**
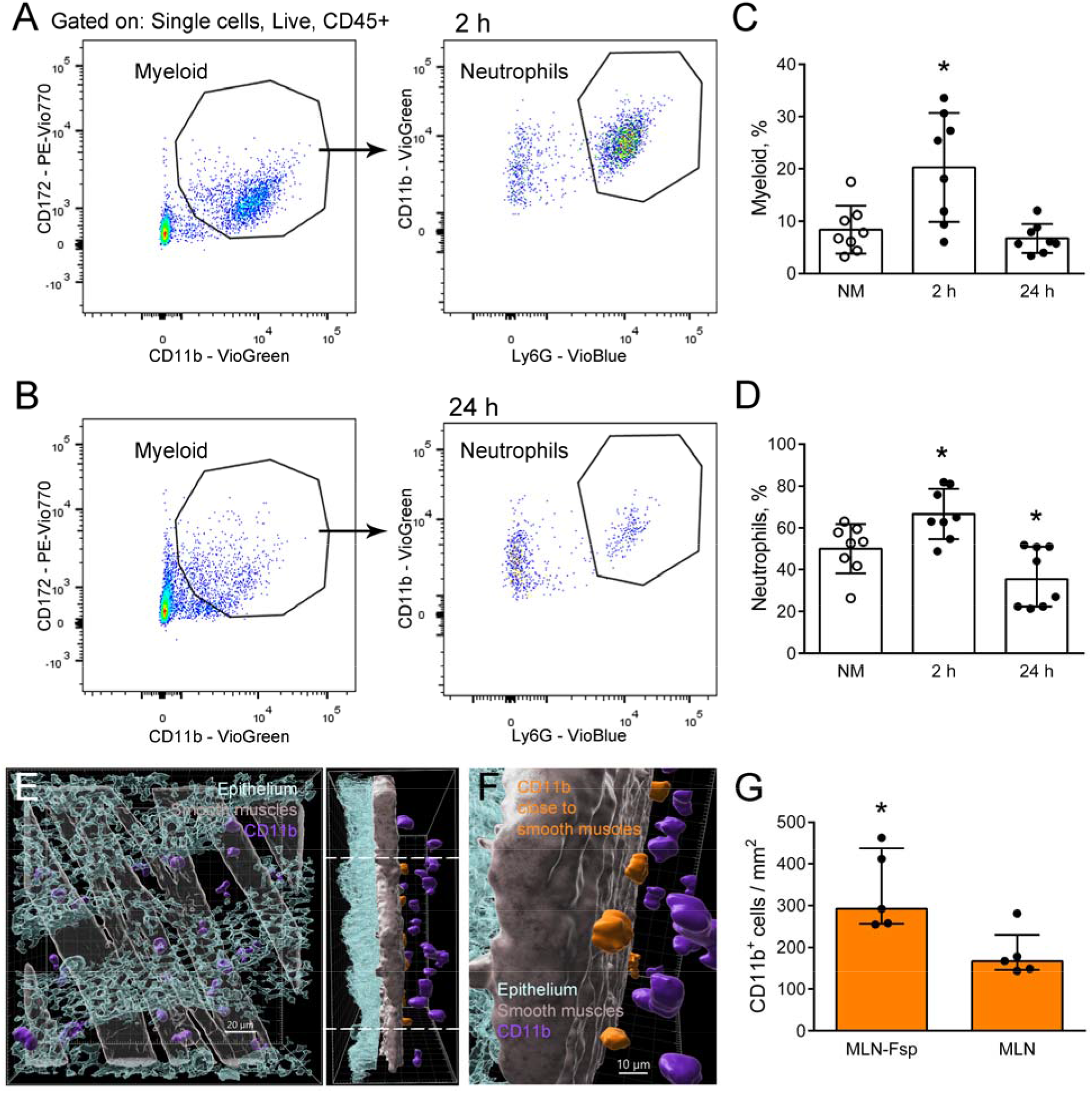
MLN-4760 and FluoSpheres oropharyngeal application induce the neutrophil retention in the conducting airway wall. A, B. Representative flow cytometry plots showing the blood myeloid cell (left plots) and neutrophil (right plots) populations of mice that received o.ph. MLN-4760 and FluoSpheres 2 h (A) or 24 h before the blood collection (B). C, D. The percentage of myeloid cells (C) and neutrophils (D) in the peripheral blood of the intact mice (NM), 2 h after the MLN-4760 and FluoSpheres o.ph. application (MLN-Fsp 2h) and 24 h after the MLN-4760 and FluoSpheres o.ph. application (MLN-Fsp 24h). The data are pooled from two independent experiments, the total number of mice n=8, and shown as median and i.q.r.; Significant difference between indicated group and intact mice *: p ≤ 0.05. E. Representative image of the conducting airway region of mice 24 h after the MLN-4760 and FluoSpheres application resented as frontal (left image) and lateral (right image) views, showing CD11b^+^ cells (violet) in the submucosa, beneath the smooth muscles (gray, transparent on the left image); the epithelium (cyan, transparent). The cells, the smooth muscle layer, and the epithelium are shown via surface rendering. FluoSpheres are not shown. The frontal (left image) and lateral (right image) projections are presented. The neutrophils located in close proximity to the smooth muscles are indicated in orange on the lateral projection. Scale bar 40 µm. F. Enlarged and rotated image of the region between the dotted lines (E, right image) showing proximal to the smooth muscles neutrophils (orange) and distal ones (violet). G. Quantitative analysis of the neutrophils proximal to the smooth muscles for mice that received MLN-4760 together with FluoSpheres and MLN-4760 alone. The data are shown as median and i.q.r., n=5 mice. Significant difference between the indicated group and the intact mice was detected using Mann– Whitney test *: p ≤ 0.05.

Thus, in the presence of nanoparticles, activated by the ACE2 inhibition neutrophils are retained in the airways.

### ACE2 blocking without RBD application is not potent to induce the neutrophil migration through the conducting airway wall to the site of the FluoSphere location

Instead of a measurable CD11b^+^ cell recruitment to the conducting airway mucosa in response to the application of MLN-4760 together with FluoSpheres, solitary CD11b^+^ cells were detected in the luminal side of the epithelium (Fig. 5A). Simultaneous application of MLN-4760, FluoSpheres, and RBD induced the neutrophil migration through the epithelium to the airway lumen (Fig. 5B). To compare the effects of the MLN-4760 applications together with FluoSpheres (in PBS) and MLN-4760 together with FluoSpheres in RBD solution on the CD11b^+^ cell recruitment, we performed a quantitative analysis of CD11b^+^ cell numbers in the conducting airway mucosa and particularly in the luminal side of the conducting airway epithelium (Fig. 5C-E). CD11b^+^ cells in the submucosa (Fig. 5C, orange) were excluded from the quantitation. The analysis showed that the number of the CD11b^+^ cells did not significantly differ in the conducting airway mucosa of mice that received FluoSpheres with or without RBD (Fig. 5D). In contrast, the number of CD11b^+^ cells in the luminal side of the epithelium was significantly higher in the presence of RBD than in mice that received MLN-4760 together with FluoSpheres without RBD (Fig. 5E). Like in the case of the application of FluoSpheres suspended in RBD solution (without MLN-4750) (Fig. 1B-D), in the conducting airways of mice that received MLN-4760 together with FluoSpheres in RBD solution, CD11b^+^ cells on the luminal side of the epithelium ingested FluoSpheres (Fig. 5F,G). Moreover, in the presence of MLN-4760 and RBD, CD11b^+^ cells contacted CD169^+^ cells that internalized FluoSpheres (Fig. 5F, arrows, G).

**Figure 5.**
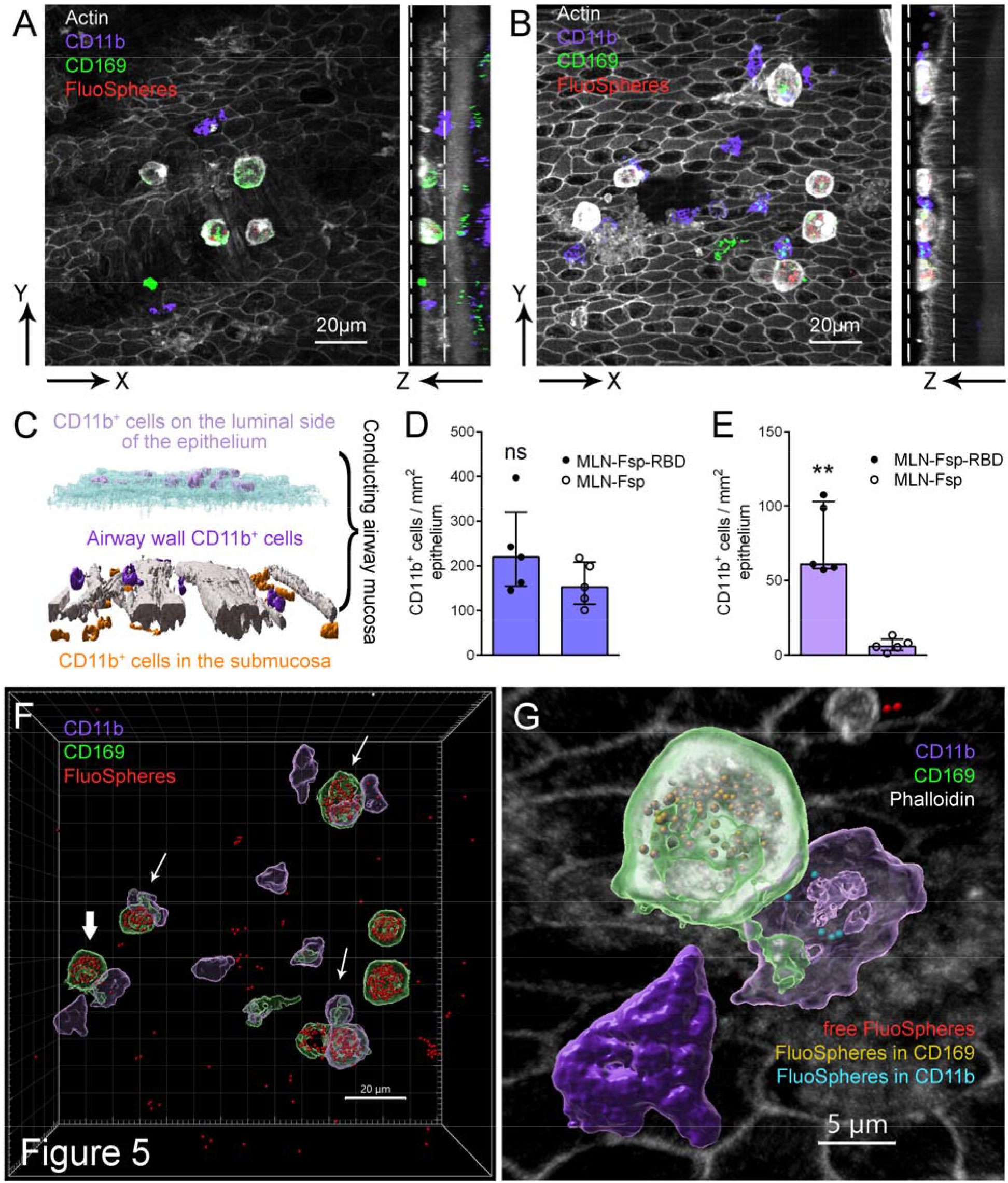
RBD contributes to the neutrophil-macrophage-FluoSphere interactions in the luminal side of the conducting airway epithelium. A, B. Representative images of the regions of the luminal side of the conducting airway epithelium of mice 24 h after the o.ph. application of FluoSpheres and MLN-4760 (A) or FluoSpheres, RBD, and MLN-4760 (B). The actin-reach filaments of the epithelium and smooth muscles (grayscale), neutrophils (violet), CD169^+^ cells (green) and FluoSpheres (red) are represented as frontal (left images) and lateral (right images) projections. Each frontal projection contains an extended view of several Z-sections approximately indicated by the dotted lines on the frontal projections. Scale bar 40 µm. C. Principle of the quantitative analysis of the conducting airway mucosa neutrophils. The image shown in (B) is presented via surface rendering: smooth muscles (white), the epithelium (cyan), CD11b^+^ cells in the submucosa (orange), CD11b^+^ cells in the airway wall (violet), and CD11b^+^ cells in the luminal side of the epithelium (light violet). D, E. The number of CD11b^+^ cells in the conducting airway mucosa (D) or on the luminal side of the conducting airway epithelium (E) of mice 24 h after the o.ph. application of FluoSpheres and MLN-4760 (MLN-Fsp) or FluoSpheres in RBD solution and MLN-4760 (MLN-Fsp-RBD). The data are shown as median and i.q.r., n=5 mice. Significant difference between the indicated groups was detected using Mann–Whitney test *: p ≤ 0.05, ns: not significant. F. The image demonstrated in (B) showing via surface rendering of the neutrophils on the luminal side of the epithelium (CD11b, violet), CD169^+^ cells (green), and FluoSpheres (red). The radius scaling was used to enlarge the FluoSphere surface to improve visual perception. The arrows indicate contacts between CD169^+^ and CD11b^+^ cells. Scale bar 20 µm. G. Enlarged image of CD11b^+^ cells (violet) and CD169^+^ cell (green) indicated on (F) with a bold arrow. FluoSpheres outside the cells (red), inside CD169^+^ cell (yellow) and inside CD11b^+^ cell (cyan). The cells and FluoSpheres are presented via surface rendering, the epithelium (grayscale) via volume rendering. The radius scaling was used to enlarge the FluoSphere surface to improve visual perception. The surfaces of the cells that internalized FluoSpheres are transparent to make FluoSpheres inside the cells visible. Scale bar 5 µm.

In sum, the airway ACE2 blocking induces the CD11b^+^ cell (including neutrophils) recruitment to the conducting airway mucosa, the nanoparticles (in our case, FluoSpheres) contribute to these CD11b^+^ cells retention in the conducting airway mucosa, and RBD triggers the neutrophil migration to the luminal side of the epithelium. On the luminal side of the epithelium, CD11b^+^ cells ingest free FluoSpheres and interact with CD169^+^ cells that have already internalized FluoSpheres.

## 3.2.6. Discussion

In the present study, we have shown that 100 nm particles reaching conducting airways undergo internalization by CD169^+^ actin-rich cells. The role of CD169^+^ monocytes and macrophages in SARS-CoV-2-induced infection is a widely discussed question (Abassi et al., 2020; Merad and Martin, 2020). The elevation of CD169^+^ activated monocytes in the periphery blood of patients was detected early upon SARS-CoV-2 infection (Chevrier et al., 2021). The implication of CD169^+^ macrophage populations, particularly lymph node subcapsular and splenic marginal zone macrophages, in SARS-CoV-2 particles internalization was reported, particularly, SARS-CoV-2 nucleoprotein was detected in such macrophages in the histological samples of the patients who died from Covid-19 (Feng et al., 2020). Further, it was shown that SARS-CoV-2 efficiently infects human monocytes and macrophages (Boumaza et al., 2021). The present study demonstrated the ability of CD169^+^ conducting airway cells to internalize 100 nm particles 24 h after the application, which supports the evidence of the contribution of CD169^+^ macrophages in the airways to internalization of virus particles, including SARS-CoV-2 particles. Although macrophages and monocytes express ACE2 and therefore can specifically interact with SARS-CoV-2 (Abassi et al., 2020), the internalization of inert FluoSpheres by CD169^+^ cells allows suggesting that virus particles can be ingested passively. However, it is still unclear if such internalization can prevent virus replication. Nevertheless, the ability to terminate SARS-CoV replication by human peripheral blood macrophages was demonstrated (Yilla et al., 2005).

Despite the data showing that nanoparticles can activate the neutrophil-mediated response (Keshavan et al., 2019), the effect depends on the experimental design and the intrinsic nanoparticle properties. Here, we have shown that inert 100 nm particles in concentrations of 1 × 10^8^ are not potent to induce the neutrophil recruitment to the conducting airway mucosa. Using initially anti-Ly6G antibodies that specifically label mouse neutrophils (Boivin et al., 2020) and then anti-CD11b antibodies (suitable for 3D immunohistochemistry) that label all phagocytic cells (including neutrophils), we were not able to identify measurable neutrophil recruitment to the conducting airway mucosa in response to inert nanoparticles. Most probably, inert particles undergo internalization by resident phagocytes (such as CD169^+^ cells) and further elimination from the bronchial branches by digestion or mucociliary clearance (Bustamante-Marin and Ostrowski, 2017).

Since neutrophils were shown to be involved in severe COVID-19 pathogenesis and associated with SARS-CoV-2-induced complications (Barnes et al., 2020; Silvin et al., 2020), we aimed to identify the factors that contribute to the neutrophil recruitment to the conducting airway mucosa in response to the SARS-CoV-2 virus particles. Considering the importance of the ACE2–RBD interaction, we checked the effect of the ACE2 blocking (by application of MLN-4760) on the neutrophil migration. Blocking or shedding of ACE2 due to the interaction with RBD of SARS-CoV-2 leads to an increase of the AngII level (Rossi et al., 2020). Some studies in humans demonstrate that AngII interaction with endothelial Angiotensin II Receptor Type 1 stimulates the elevation of leukocyte attractants, including neutrophil attractant IL-8 (Bernstein et al., 2018).

Besides, the increased AngII level induces the ATP release by erythrocytes (Zhao et al., 2019). The elevation of extracellular ATP at the site of inflammation, in turn, stimulates the neutrophil migration (Karmakar et al., 2016). In the present study, we have shown that the ACE2 inhibition both systemic and local (in the airways) induces the neutrophil recruitment from the bone marrow to the bloodstream. In the case of the airway ACE2 inhibition, neutrophils also migrated from the bloodstream further to the lungs. Moreover, in the presence of nanoparticles, neutrophils were retained in the airways. However, nanoparticles were located in the luminal side of the epithelium, while neutrophils were located in the conducting airway wall; they did not migrate through the epithelial barrier in response to inert nanoparticles.

Massive neutrophil migration to the site of the pathogen entrance or tissue damage is well described, but the molecular mechanisms regulating the transepithelial migration are not completely understood (Lin and Fessler, 2021). In the case of COVID-19, the interaction of neutrophils with the SARS-CoV-2 virus particles was shown to contribute to the NET formation (Veras et al., 2020). Here, we have shown that 24 h after the oropharyngeal application, nanoparticles were located in the luminal side of the epithelium, while neutrophils migrated from the vessels and submucosal compartment. The transepithelial neutrophil migration was necessary for the neutrophil-nanoparticle contact, however inert nanoparticles even in conditions of the ACE2 blocking did not stimulate the transepithelial neutrophil migration. RBD appears to be essential for the neutrophil migration through the airway epithelial barrier. Although BSA did not stimulate the neutrophil attraction to the airway lumen, the ability to trigger the neutrophil transepithelial migration can be a non-exclusive attribute of RBD. Moreover, it is still unclear which secondary messengers are involved in the RBD-mediated triggering of the neutrophil transepithelial migration. This question is important to determine the mechanisms of the acute airway inflammatory disorders, including virus-induced inflammation, and to develop strategies to suppress such inflammation.

Our results demonstrate that the ACE2 inhibition together with the RBD-mediated neutrophil attraction results in sufficient neutrophil recruitment to the airways and migration to the luminal side of the epithelium, which spatially allows neutrophil-particle interactions. Besides, the synergetic effect of the ACE2 inhibition and the RBD-mediated neutrophil activation results in the neutrophil-macrophage contacts in the luminal side of the conducting airway epithelium. The neutrophil-macrophage contacts contribute to inflammation resolution and tissue repair (Bouchery and Harris, 2019; Marwick et al., 2018). Simultaneously, neutrophils swarming can uncover resident tissue macrophages-mediated cloaking of pathogens or tissue lesions and switch the tolerance to inflammation (Uderhardt et al., 2019). Although the nature of the neutrophil-macrophage contacts reported in the present study is still unclear, the observation may suggest that in the presence of RBD, CD169^+^ cells can lose control of the nanoparticle dissemination and neutrophil recruitment blocking.

Thus, here we report the approach that allows mimicking the RBD–ACE2 interaction in mice and simulating virus particle loading. Using this approach, we have demonstrated that the ACE2 inhibition activates the neutrophil migration from the bone marrow to the bloodstream, nanoparticles facilitate the neutrophil retention in the lungs, and RBD contributes to the transepithelial neutrophil migration and nanoparticle internalization.

## 3.2.7. Materials and Methods

### Animals and ethics statement

C57BL/6 mice were purchased from Pushchino Animal Facility of the Shemyakin and Ovchinnikov Institute of Bioorganic Chemistry, Russian Academy of Sciences; female (18-20 g, 8-12 weeks). All animal experiments were performed in concordance with the Guide for the Care and Use of Laboratory Animals under a protocol approved by the Institutional Animal Care and Use Committee at Shemyakin and Ovchinnikov Institute of Bioorganic Chemistry Russian Academy of Sciences (protocol numbers 302/2020, 319/2021). The animals were given standard food and tap water ad libitum and housed under regular 12-h dark:light cycles at 22°C.

### Fluorescent particles

Fluorescent particles FluoSpheres Carboxylate-Modified Microspheres, 100 nm, red fluorescent (580/605) (ThermoFisher, F8801) were used in the study. The particle concentration was estimated in accordance with the manufacturer recommendations; the particles were dissolved to the concentration of 1.7 × 10^8^ before the application to mice. The particles were dissolved in PBS (Paneco, Russia) or in 0.1% solution of RBD (Hytest, Russia, 8COV1) or in 0.1% bovine serum albumin (BSA) (Sigma-Aldrich).

### ACE2 inhibitor application

Mice received an i.p. injection of the ACE2 inhibitor MLN-4760 (Merck, 5.30616.0001). MLN-4760 was dissolved in PBS and injected to the mice in a total volume of 200 µL in dose 0.04 mg/mouse. Control mice received the i.p. injection of PBS. In the other experiments, mice were anesthetized by inhalation of 0.5-3 % isoflurane (Baxter, Guayama, Puerto Rico) and then received the o.ph. application of MLN-4760 in dose 0.04 mg per mouse per inhalation in total volume of 50 µL per mouse per inhalation. Control mice received the o.ph. application of PBS.

### Particles application

Mice were anesthetized by inhalation of 0.5-3 % isoflurane, and a 50-μL droplet containing particles was applied to the oropharyngeal cavity of each mouse. Control mice received PBS or 0.1% RBD, or 0.1% BSA.

### Blood collection

The peripheral blood was collected from the tail vein. The blood was collected to the 1.5 mL test-tube with 50 µL heparin (VelPharm, Russia) and transferred to the 10 mL of the hemolysis buffer (155 mM NH4Cl (Reachem, Russia); 0.1 mM Na_2_EDTA (Sigma-Aldrich); 10 mM NaHCO_3_ (PanReac Applichem); pH 7.3, stored at +4°C, and warmed to RT before use) in 50 mL tube. After 5 min at RT, 20 mL of PBS was added, and the tube was centrifuged at 380 g 5 min. The supernatant was replaced with 5 mL of the hemolysis buffer. After 5 min at RT, cells were centrifuged at 380 g 5 min and transferred to 500 µL of the cytometry buffer (1% BSA 2 mM EDTA). Then cells were centrifuged using CV 1500 (Biosan, Latvia) for 10 min. The cell pellet was dissolved in 30 µL of the cytometry buffer and placed to the 96 well plate.

### Blood cell analysis by flow cytometry

For the flow cytometry analysis, we used the strategy for the peripheral blood myeloid cell detection, which was recommended by Liu et al. (Liu et al., 2020). The samples were preincubated with anti-mouse CD16/CD32 (Miltenyi Biotec, 130-092-574) for 15 min. Then the following antibodies (all from Miltenyi Biotec) were used: anti-mouse Ly6G–VioBlue (130-119-902), anti-mouse CD11b–VioGreen (130-113-811), anti-mouse FcεR1–PE (130-118-896), anti-mouse CD172–PE-Vio770 (130-123-154), anti-mouse CD45–APC-Vio770 (130-110-800). The antibodies were used in dilution 1:30. The samples were incubated for 30 min and washed twice with PBS. SytoxGreen (Invitrogen, S34859) was added (in dilution 1:1000000) 5 min before the acquisition. The measurement was performed using a MACSQuant Analyzer 10 (Miltenyi Biotec).

### Whole-mount conducting airway specimen preparation and staining

The animals were euthanized and their lungs were fixed with 2% paraformaldehyde without inflation and stored at 4 °C overnight. The main bronchi from the lung lobes (left and right inferior) were dissected. The main bronchi from the lung lobes were dissected. The left, and right inferior lobes were then used for the quantitative analysis, the left middle or post-caval lobes were used for isotype-control staining (Fig. S4A, B). The airway specimens were then washed with PBS, permeabilized with 0.3% Triton X-100, and blocked with 1% BSA and 4% normal goat serum and/or normal donkey serum (Jackson Immuno Research, 005-000-121) or with 1% BSA. The following antibodies and dilutions were used: anti-mouse CD11b-APC (BioLegend, 101212, dilution 1:50), anti-mouse CD169-AlexaFluor488 (BioLegend, 142419, dilution 1:50), anti-mouse Ly6G-AlexaFluor647 (BioLegend, 127609, dilution 1:50). If indicated, anti-mouse CD326-AlexaFluor594 (BioLegend, 118222, dilution 1:100) were used for labeling the epithelium and unconjugated goat anti-mouse CD31 (R&D Systems, AF3628, dilution 1:100) in combination with secondary donkey anti-goat-AlexaFluor555 (ThermoFisher, A-21434, dilution 1:300) were used for labeling the blood vessels. For isotype-control staining APC-conjugated Rat IgG2b, κ (BioLegend, 400611) and Rat IgG2a, κ (BioLegend, 400525) The specimens were incubated with antibodies overnight then washed with 0.3% Triton X-100 in PBS and stained against actin by incubation for 1 h with Phalloidin-Atto425 (Sigma-Aldrich, 66939, dilution 1:50). If indicated, nuclei were stained with Hoechst (PanEco, dilution 1:1000) and subsequent washing with PBS. All samples were covered with Prolong Gold mounting medium (Thermo Fisher, P36930).

### Confocal Laser-Scanning Microscopy

An inverted confocal LSM780 microscope (Zeiss, Jena, Germany) was used in all experiments with either a 10× (NA = 0.3), 40× (NA = 1.4, water immersive), or a 100× (NA = 1.46, oil immersive) objective. Excitation at 405, 488, 561, and 633 nm was used to visualize Atto425, Alexa Fluor 488, Alexa Fluor 555/594, and APC, respectively. Emission was measured in CLSM λ-mode using a 34-channel QUASAR detector (Zeiss) set to a 405–695 nm range. For quantitative analysis, the images were captured as 2 × 2 tile grids at the same regions of each specimen using the 40× objective, with an individual xyz tile size of 354 µm × 354 µm × 30 µm. Higher magnification images were acquired in z-stacks at the region of interest using the 100× objective. The images were acquired as three-dimensional stacks. Spectral unmixing was performed using ZEN 2012 SP5 software (Zeiss). The images were created using Imaris version 9.8 software (Oxford Instruments). Cells and FluoSpheres were presented either as maximum intensity projections or as surfaces that were created based on the maximum intensity projections. The FluoSpheres surfaces were enlarged using the radius-scaling function to improve visual perception. Finally, the images were processed using Adobe Photoshop CS version 5 (Adobe Systems, Mountain View, CA).

### Quantitative Image Analysis

The image stacks were analyzed using Imaris. Based on maximum intensity projections, CD11b^+^ cells were processed via three-dimensional surface rendering of the APC channel, using several thresholds, as previously described (Veres et al., 2009). CD11b^+^ cell surfaces were quantified automatically; the inspection of the results was made each time manually. The images of at least two proximate to trachea regions from each specimen were acquired (Shevchenko et al., 2013).

### Statistical Analysis

The data are presented as the scattered dot plots with the median and interquartile range (i.q.r.) for at least 4 mice per group. The differences between two groups were analyzed with the Mann– Whitney test using GraphPad Prism software (GraphPad Software, San Diego, CA). A p-value less than 0.05 was considered statistically significant.

## 3.2.8. Acknowledgments

The authors thank for editing this manuscript the Head of Foreign Languages Department of Moscow Institute of Physics and Technology, associate professor of MIPT Dr. Elena Bazanova.

## 3.2.11. Data availability

## 3.2.12. Author contributions statement

IO, VB and MS conceptualization; JV, IO and MS methodology; EB, JV, AB, YZ and MS investigation; MS original draft preparation; AB and MS review and editing; AS and VB supervision; MS project administration and funding acquisition.

## References

Abassi, Z., Knaney, Y., Karram, T. and Heyman, S. N. (2020). The Lung Macrophage in SARS-CoV-2 Infection: A Friend or a Foe? Frontiers in Immunology 11, 1312.

Bao, L., Deng, W., Huang, B., Gao, H., Liu, J., Ren, L., Wei, Q., Yu, P., Xu, Y., Qi, F., et al. (2020). The pathogenicity of SARS-CoV-2 in hACE2 transgenic mice. Nature 583, 830– 833.

Barnes, B. J., Adrover, J. M., Baxter-Stoltzfus, A., Borczuk, A., Cools-Lartigue, J., Crawford, J. M., Daßler-Plenker, J., Guerci, P., Huynh, C., Knight, J. S., et al. (2020). Targeting potential drivers of COVID-19: Neutrophil extracellular traps. Journal of Experimental Medicine 217, e20200652.

Bar-On, Y. M., Flamholz, A., Phillips, R. and Milo, R. (2020). SARS-CoV-2 (COVID-19) by the numbers. eLife 9, e57309.

Bernstein, K. E., Khan, Z., Giani, J. F., Cao, D. Y., Bernstein, E. A. and Shen, X. Z. (2018). Angiotensin-converting enzyme in innate and adaptive immunity. Nature Reviews Nephrology 14, 325–336.

Bogorodskiy, A. O., Bolkhovitina, E. L., Gensch, T., Troyanova, N. I., Mishin, A. v., Okhrimenko, I. S., Braun, A., Spies, E., Gordeliy, V. I., Sapozhnikov, A. M., et al. (2020). Murine Intraepithelial Dendritic Cells Interact With Phagocytic Cells During Aspergillus fumigatus-Induced Inflammation. Frontiers in Immunology 11, 298.

Boivin, G., Faget, J., Ancey, P.-B., Gkasti, A., Mussard, J., Engblom, C., Pfirschke, C., Contat, C., Pascual, J., Vazquez, J., et al. (2020). Durable and controlled depletion of neutrophils in mice. Nature Communications 11, 2762.

Bouchery, T. and Harris, N. (2019). Neutrophil–macrophage cooperation and its impact on tissue repair. Immunology & Cell Biology 97, 289–298.

Boumaza, A., Gay, L., Mezouar, S., Bestion, E., Diallo, A. B., Michel, M., Desnues, B., Raoult, D., la Scola, B., Halfon, P., et al. (2021). Monocytes and Macrophages, Targets of Severe Acute Respiratory Syndrome Coronavirus 2: The Clue for Coronavirus Disease 2019 Immunoparalysis. The Journal of Infectious Diseases 224, 395–406.

Bustamante-Marin, X. M. and Ostrowski, L. E. (2017). Cilia and mucociliary clearance. Cold Spring Harbor Perspectives in Biology 9, a028241.

Chevrier, S., Zurbuchen, Y., Cervia, C., Adamo, S., Raeber, M. E., de Souza, N., Sivapatham, S., Jacobs, A., Bachli, E., Rudiger, A., et al. (2021). A distinct innate immune signature marks progression from mild to severe COVID-19. Cell Reports Medicine 2, 100166.

Cicko, S., Köhler, T. C., Ayata, C. K., Müller, T., Ehrat, N., Meyer, A., Hossfeld, M., Zech, A., di Virgilio, F. and Idzko, M. (2018). Extracellular ATP is a danger signal activating P2×7 receptor in a LPS mediated inflammation (ARDS/ALI). Oncotarget 9, 30635– 30648.

Feng, Z., Diao, B., Wang, R., Wang, G., Wang, C., Tan, Y., Liu, L., Wang, C., Liu, Y., Liu, Y., et al. (2020). The Novel Severe Acute Respiratory Syndrome Coronavirus 2 (SARS-CoV-2) Directly Decimates Human Spleens and Lymph Nodes. medRxiv doi:10.1101/2020.03.27.20045427

Gu, H., Chen, Q., Yang, G., He, L., Fan, H., Deng, Y.-Q., Wang, Y., Teng, Y., Zhao, Z., Cui, Y., et al. (2020). Adaptation of SARS-CoV-2 in BALB/c mice for testing vaccine efficacy. Science 369, 1603–1607.

Gusarova, G. A., Das, S. R., Islam, M. N., Westphalen, K., Jin, G., Shmarakov, I. O., Li, L., Bhattacharya, S. and Bhattacharya, J. (2021). Actin fence therapy with exogenous V12Rac1 protects against acute lung injury. JCI Insight 6, e135753.

Hazeldine, J. and Lord, J. M. (2021). Neutrophils and COVID-19: Active Participants and Rational Therapeutic Targets. Front. Immunol. 12, 680134.

Karmakar, M., Katsnelson, M. A., Dubyak, G. R. and Pearlman, E. (2016). Neutrophil P2×7 receptors mediate NLRP3 inflammasome-dependent IL-1β secretion in response to ATP. Nature Communications 7, 10555.

Keshavan, S., Calligari, P., Stella, L., Fusco, L., Delogu, L. G. and Fadeel, B. (2019). Nano-bio interactions: a neutrophil-centric view. Cell Death & Disease 10, 569.

Lin, W.-C. and Fessler, M. B. (2021). Regulatory mechanisms of neutrophil migration from the circulation to the airspace. Cellular and Molecular Life Sciences 78, 4095–4124.

Liu, Z., Gu, Y., Shin, A., Zhang, S. and Ginhoux, F. (2020). Analysis of Myeloid Cells in Mouse Tissues with Flow Cytometry. STAR Protocols 1, 100029.

Marwick, J. A., Mills, R., Kay, O., Michail, K., Stephen, J., Rossi, A. G., Dransfield, I. and Hirani, N. (2018). Neutrophils induce macrophage anti-inflammatory reprogramming by suppressing NF-κB activation. Cell Death & Disease 9, 665.

Merad, M. and Martin, J. C. (2020). Pathological inflammation in patients with COVID-19: a key role for monocytes and macrophages. Nature Reviews Immunology 20, 355–362.

Middleton, E. A., He, X. Y., Denorme, F., Campbell, R. A., Ng, D., Salvatore, S. P., Mostyka, M., Baxter-Stoltzfus, A., Borczuk, A. C., Loda, M., et al. (2020). Neutrophil extracellular traps contribute to immunothrombosis in COVID-19 acute respiratory distress syndrome. Blood 136, 1169–1179.

Mutua, V. and Gershwin, L. J. (2021). A Review of Neutrophil Extracellular Traps (NETs) in Disease: Potential Anti-NETs Therapeutics. Clinical Reviews in Allergy and Immunology 61, 194–211.

Rossi, G. P., Sanga, V. and Barton, M. (2020). Potential harmful effects of discontinuing ACE-inhibitors and ARBs in COVID-19 patients. eLife 9, e57278.

Silvin, A., Chapuis, N., Dunsmore, G., Goubet, A.-G., Dubuisson, A., Derosa, L., Almire, C., Hénon, C., Kosmider, O., Droin, N., et al. (2020). Elevated Calprotectin and Abnormal Myeloid Cell Subsets Discriminate Severe from Mild COVID-19. Cell 182, 1401-1418.e18.

Shevchenko, M. A., Bolkhovitina, E. L., Servuli, E. A. and Sapozhnikov, A. M. (2013). Elimination of Aspergillus fumigatus conidia from the airways of mice with allergic airway inflammation. Respir. Res. 14, 78.

Uderhardt, S., Martins, A. J., Tsang, J. S., Lämmermann, T. and Germain, R. N. (2019). Resident Macrophages Cloak Tissue Microlesions to Prevent Neutrophil-Driven Inflammatory Damage. Cell 177, 541-555.e17.

Veras, F. P., Pontelli, M. C., Silva, C. M., Toller-Kawahisa, J. E., de Lima, M., Nascimento, D. C., Schneider, A. H., Caetité, D., Tavares, L. A., Paiva, I. M., et al. (2020). SARS-CoV-2–triggered neutrophil extracellular traps mediate COVID-19 pathology. Journal of Experimental Medicine 217,e20201129.

Veres, T. Z., Shevchenko, M., Krasteva, G., Spies, E., Prenzler, F., Rochlitzer, S., Tschernig, T., Krug, N., Kummer, W. and Braun, A. (2009). Dendritic cell-nerve clusters are sites of T cell proliferation in allergic airway inflammation. American Journal of Pathology 174, 808–17.

Wang, Q., Zhang, Y., Wu, L., Niu, S., Song, C., Zhang, Z., Lu, G., Qiao, C., Hu, Y., Yuen, K. Y., et al. (2020). Structural and Functional Basis of SARS-CoV-2 Entry by Using Human ACE2. Cell 181, 894-904.e9.

Winkler, E. S., Chen, R. E., Alam, F., Yildiz, S., Case, J. B., Uccellini, M. B., Holtzman, M. J., Garcia-Sastre, A., Schotsaert, M. and Diamond, M. S. (2022). SARS-CoV-2 Causes Lung Infection without Severe Disease in Human ACE2 Knock-In Mice. Journal of Virology 96, e0151121.

Yang, S. C., Tsai, Y. F., Pan, Y. L. and Hwang, T. L. (2021). Understanding the role of neutrophils in acute respiratory distress syndrome. Biomed. J. 44, 439–446.

Ye, M., Wysocki, J., Gonzalez-Pacheco, F. R., Salem, M., Evora, K., Garcia-Halpin, L., Poglitsch, M., Schuster, M. and Batlle, D. (2012). Murine Recombinant Angiotensin-Converting Enzyme 2. Hypertension 60, 730–740.

Yilla, M., Harcourt, B. H., Hickman, C. J., McGrew, M., Tamin, A., Goldsmith, C. S., Bellini, W. J. and Anderson, L. J. (2005). SARS-coronavirus replication in human peripheral monocytes/macrophages. Virus Research 107, 93–101.

Zhao, T. v., Li, Y., Liu, X., Xia, S., Shi, P., Li, L., Chen, Z., Yin, C., Eriguchi, M., Chen, Y., et al. (2019). ATP release drives heightened immune responses associated with hypertension. Science Immunology 4, eaau6426.

Zhou, P., Yang, X.-L., Wang, X.-G., Hu, B., Zhang, L., Zhang, W., Si, H.-R., Zhu, Y., Li, B., Huang, C.-L., et al. (2020). A pneumonia outbreak associated with a new coronavirus of probable bat origin. Nature 579, 270–273.

Zuo, Y., Yalavarthi, S., Shi, H., Gockman, K., Zuo, M., Madison, J. A., Blair, C. N., Weber, A., Barnes, B. J., Egeblad, M., et al. (2020). Neutrophil extracellular traps in COVID-19. JCI Insight 4, e138999.

